# Correlations of 276 missense variants of *PSEN1, PSEN2*, and *APP* on the production of Aβ peptides against variant effect predictors and biophysical structures

**DOI:** 10.64898/2026.02.10.705105

**Authors:** Jiyeon Song, Serina Yan, Sophia Liu, Adhvaith Sridhar, Kijung Sung, Joshua Pillai, Chengbiao Wu

## Abstract

**INTRODUCTION:** Early-onset Alzheimer’s disease (EOAD) is mostly caused by mutations in Presenilin-1,2 (*PSEN1, PSEN2)*, and amyloid precursor protein (*APP)* genes. Both genetic and experimental evidence has supported the PSEN-APP amyloid hypothesis as the leading pathogenesis of AD. Thus far, 276 missense variants from *PSEN1,2*, and *APP* have been identified. Herein, we report the development of algorithm platforms for predicting the degrees of pathogenic potentials of these various missense mutations in AD.

**METHODS:** We performed extensive pairwise correlations between 276 *in vitro* functional assays of missense variants from *PSEN1, PSEN2*, and *APP* and clinical data among EOAD patients against 37 variant effect predictors (VEPs). Furthermore, we used the structural models of all three proteins to refine understanding of the molecular basis of missense variants in dominant negative (DN) or loss-of-function (LoF) effects.

**RESULTS:** We found that VEPs were consistently correlated with age of onset (AAO) among EOAD patients and the Aβ42/Aβ40 ratio biomarker. This study also identified discrepancies in the predictor variable leading to an increase in the Aβ42/Aβ40 ratio; The increased Aβ42/Aβ40 ratio was due to decreased Aβ40 levels by Sun et al., (2016), while Petit et al., (2022) argued that increased Aβ42 level was the culprit. We also identified that with increased predicted pathogenicity, there were decreased Aβ38 levels. Conversely, this study found no direct correlations with Aβ37 and Aβ43. Lastly, structural studies show characteristics of either DN and LoF mechanisms, as there were numerous variants located in buried hydrophobic domains and those on charged surfaces of the proteins.

**DISCUSSION:** Currently, a major limitation in the field is the lack of reliable approaches for predicting the pathogenicity of variants in EOAD. We show that VEPs are promising in deciphering the effects of variants in *PSEN1, PSEN2*, and *APP* genes, especially between multiple pathogenic mutations. Our study also illuminates the need for experimental studies to identify the predictor variable causing the increased Aβ42/Aβ40 biomarker. Our biophysical studies also identify the need to characterize whether the mechanism of pathogenesis caused by DN or LoF effects is in scope of the presenilin hypothesis.

**Research in Context:**

1. **Systematic review:** The authors reviewed experimental evidence of pathogenic *PSEN1, PSEN2*, and *APP* variants in scope of the presenilin hypothesis. To our knowledge, the pathogenesis of EOAD remains incompletely understood. There are also no predictive tools/models available for informing clinical translation.
2. **Interpretation:** VEPs demonstrate strong correlations with AAO and Aβ levels from pairwise correlations with 276 missense variants. Structurally, it appears that mutations demonstrate characteristics of dominant negative and loss-of-function effects.
3. **Future Direction:** Larger studies are necessary of VEPs for realistic clinical usage.

**Highlights:**

- We performed correlations of 37 VEPs against *in vitro* data of EOAD variants.
- VEPs demonstrate potential as diagnostic tools for early onset Alzheimer’s disease.
- Pathogenic missense variants show characteristics of DN and LoF effects.

## 1. BACKGROUND

*Presenilin* genes have been identified as major genetic linkages with early onset familial Alzheimer’s disease (EOAD), they encode the catalytic subunit of γ-secretase that cleaves type I transmembrane proteins such as amyloid precursor protein (*APP*) to release Aβ peptides (Yan et al., 2024). Though the wild-type function of these genes are essential to neuronal survival, the effects of missense variants remain controversial as to whether it causes dominant negative effects or increased Aβ42/Aβ40 levels. In the landmark reporting from Sun et al., (2016), there was a lack of statistical evidence between the Aβ42/Aβ40 levels and age of onset (AAO) among AD patients. In Szaruga et al., (2016), a unifying model was postulated where mutations led to destabilization of the γ-secretase in addition to other environmental factors to form amyloidogenic Aβ. Likewise, this model has received experimental support from Petit et al., (2022), identifying trends among the entire Aβ spectrum. Petit et al., (2022) suggest that the negative data from Sun et al., (2016) is due to the experimental conditions of the cells *in vitro*. Most importantly, their study found a linear relationship between the Aβ (37□+ □38□+ □40) □/ □ (42□+ □43) ratio and AAO. In our current understanding, we still have not identified the minimum change in Aβ profiles for AAO for direct usage in therapeutic settings and genetic counseling.

Over the past decade, missense variant effect predictors (VEPs) have been developed to predict missense mutations pathogenicity based on sequential patterns, evolutionary conservation, and recently secondary structure. These *in silico* approaches have broad applicability in precision medicine applications, and have been used for numerous Mendelian disorders (Chabane et al., 2023; Krenn et al., 2025; Staklinski et al., 2023; Wang et al., 2024a; Wang et al., 2024b). In particular, AlphaMissense (AM) has shown correlation with *in vitro* functional assays, the American College of Medical Genetics and Genomics classification, and other genetic factors (Cheng et al., 2023). Previously, we performed brief pairwise correlations between CADD, AM, ESM-1b, and EVE with the 114 VUS of *PSEN1, PSEN2*, and *APP*, and found that increased Aβ42/Aβ40 levels were driven by decreased Aβ40 levels, but lacked statistical significance for individual Aβ levels (Pillai et al., 2025). The main limitation was the limited sample size of variants, predictors, and biochemical/clinical data used for analysis.

Herein, we conducted extensive correlational analyses of 37 variant effect predictors with *in vitro* functional assays of Aβ42, 40, 38, and 37 along with AAO clinical data from EOAD patients. We also performed pairwise correlations with biophysical data of these proteins against clinical and biochemical levels. This cross-sectional study is critical to providing validation of the translational capabilities of VEPs for AD, and refine our understanding of the presenilin hypothesis.

## 2. METHODS

### 2.1. In vitro functional assays, patient data, and variant effect predictors

All *in vitro* functional data for *PSEN1, PSEN2*, and *APP* variants were based on published studies. Sun et al., (2016) had assays measuring Aβ42, Aβ40 levels for 138 pathogenic *PSEN1* variants, of which included 126 missense variants. Petit et al., (2022) reported 36 pathogenic missense *PSEN1* variants and measured Aβ42, Aβ40, Aβ38, Aβ37 levels. From our previous study, we included 56 *PSEN1*, 33 *PSEN2*, and 25 *APP* variants derived from Hsu et al., (2020) and Marsh et al., (2025). The age of onset (AAO) among patients with *PSEN1* variants were directly obtained from Sun et al., (2016). Lastly, data for all VEPs was obtained from dbNSFP (Liu et al., 2011), specifically ‘rankscore’ columns.

### 2.3. Biophysical structures

There are no full-length structures of all three amyloidogenic proteins reported from the Protein Data Bank. Therefore, we used AlphaFold 3 (AF3) to create a template (Abramson et al., 2024). We superimposed Cryo-EM structures of presenilin-1 at 2.40 L (PDB: 8KCS) and presenilin-2 at 3.00 L (PDB: 7Y5X) for validation. The DSSP library calculated relative solvent accessibilities (RSA) of residues (Tien et al., 2013).

### 2.4. Statistical analyses

Statistical analyses performed in this study used GraphPad Prism (version 10.0.0). All data are shown as mean ± standard error of the mean (SEM). Unless stated otherwise, α =□0.05 and *p*-values are indicated in the corresponding figures and tables. Spearman’s correlation was calculated in the scipy library.

## RESULTS

### 3.1. Variant effect predictors correlate strongly with age of onset and Aβ42/Aβ40 levels, but not with individual Aβ levels among amyloidogenic variants

Using the aggregated functional assays data for *PSEN1, PSEN2*, and *APP* missense variants, we performed correlations against all 37 VEPs. Within each heatmap, we have provided the Spearman’s correlation value for each individual predictor. In **Figure S1-5**, we correlated each individual predictor against all outcomes, including biochemical Aβ levels and clinical data as heatmaps. In this main text, we have displayed the visualizations for the exemplary correlations of AlphaMissense (AM) to summarize the broad associations for each gene.

Beginning with the *PSEN1* variants from Sun et al., (2016), we have identified multiple trends in predictive pathogenicity and outcomes among each gene The most important predictor variable was AAO, and we found that most VEPs had a significantly negative correlation (32/37; 86.48%). In **Figure 1A**, we show the exemplary correlation with AlphaMissense (AM) that is significantly correlated albeit outliers. For the critical Aβ42/Aβ40 levels in *PSEN1* variants, the majority of algorithms had a significantly positive correlation (29/37; 78.37%) **(Figure 1B)**. Our analyses sought to further refine the cause of increased Aβ42/Aβ40 levels with individual Aβ42, Aβ40 levels **(Figure 1C-D)**. In Aβ42, more than 80% of VEPs failed to detect significant reductions with increased predicted pathogenicity (6/37; 16.22%). Similarly, most techniques lacked significance with Aβ40 levels (18/37, 48.65%). However, we identify stronger negative associations with Aβ40 **(Figure S3)**, supporting our previous hypothesis from Pillai et al., (2025) that the increased Aβ42/Aβ40 ratio is driven by a marked decrease in Aβ40 levels in *PSEN1*.

**Figure 1.**
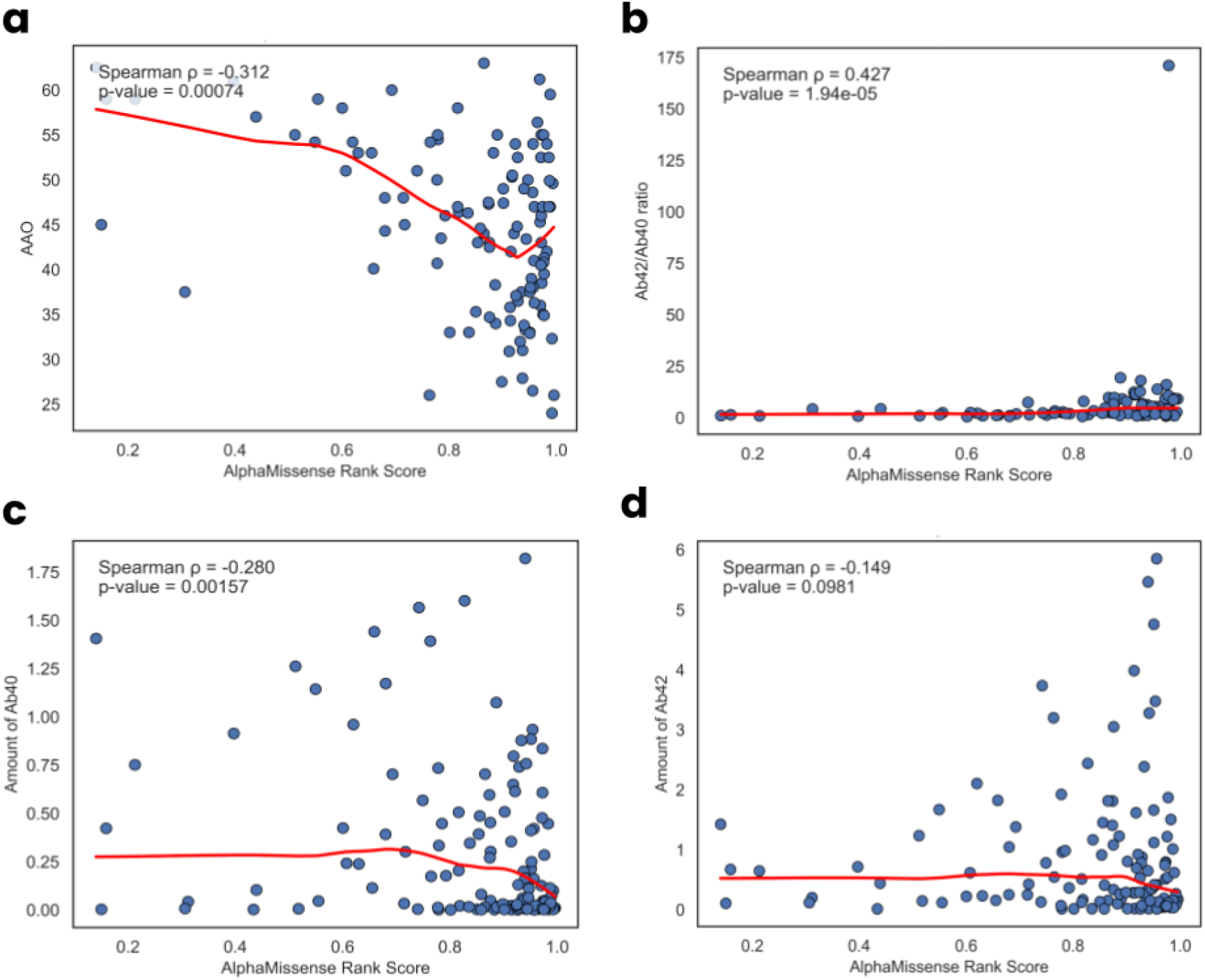
Correlations of AlphaMissense predictor with variants from Sun et al. (2016). **(a)** Correlation of AAO with AM. **(b)** Correlation between Aβ42/Aβ40 ratio and AM. **(c)** Correlation of Aβ40 and AM. **(d)** Correlation Aβ42 levels with AM.

Our analyses continued with those *PSEN1* variants from Petit et al., (2022), with emphasis on correlations with Aβ37, Aβ38, and Aβ43 **(Figure 2A-C)**. We identified no significant correlations between the VEPs and Aβ37 levels (5/37; 13.51%). More than half of all algorithms detected a significant negative correlation with Aβ38 levels (23/37; 62.16%). Likewise, only a minority of variants demonstrated positive correlations with Aβ43 levels (8/37; 21.62%). Overall, there is generally weak significance among VEPs and Aβ37, Aβ38, and Aβ43 levels among *PSEN1* variants. Next, our correlational analyses sought to validate the data from Sun et al., (2016). Surprisingly, we consistently showed no significant reductions in Aβ40 levels (2/37; 5.41%) similarly to Aβ37 levels. Even more astonishing is that more than half of all VEPs had positive correlations with Aβ42 levels (20/37; 54.05%) **(Figure 2D)**. Overall, there appears to be major discrepancies observed in the Aβ40 and Aβ42 levels between the variant samples Petit et al., (2022) and Sun et al., (2016) for the underlying mechanism for increased Aβ42/Aβ40 ratio in *PSEN1* variants and upholds the postulate proposed from Petit et al., (2022).

**Figure 2.**
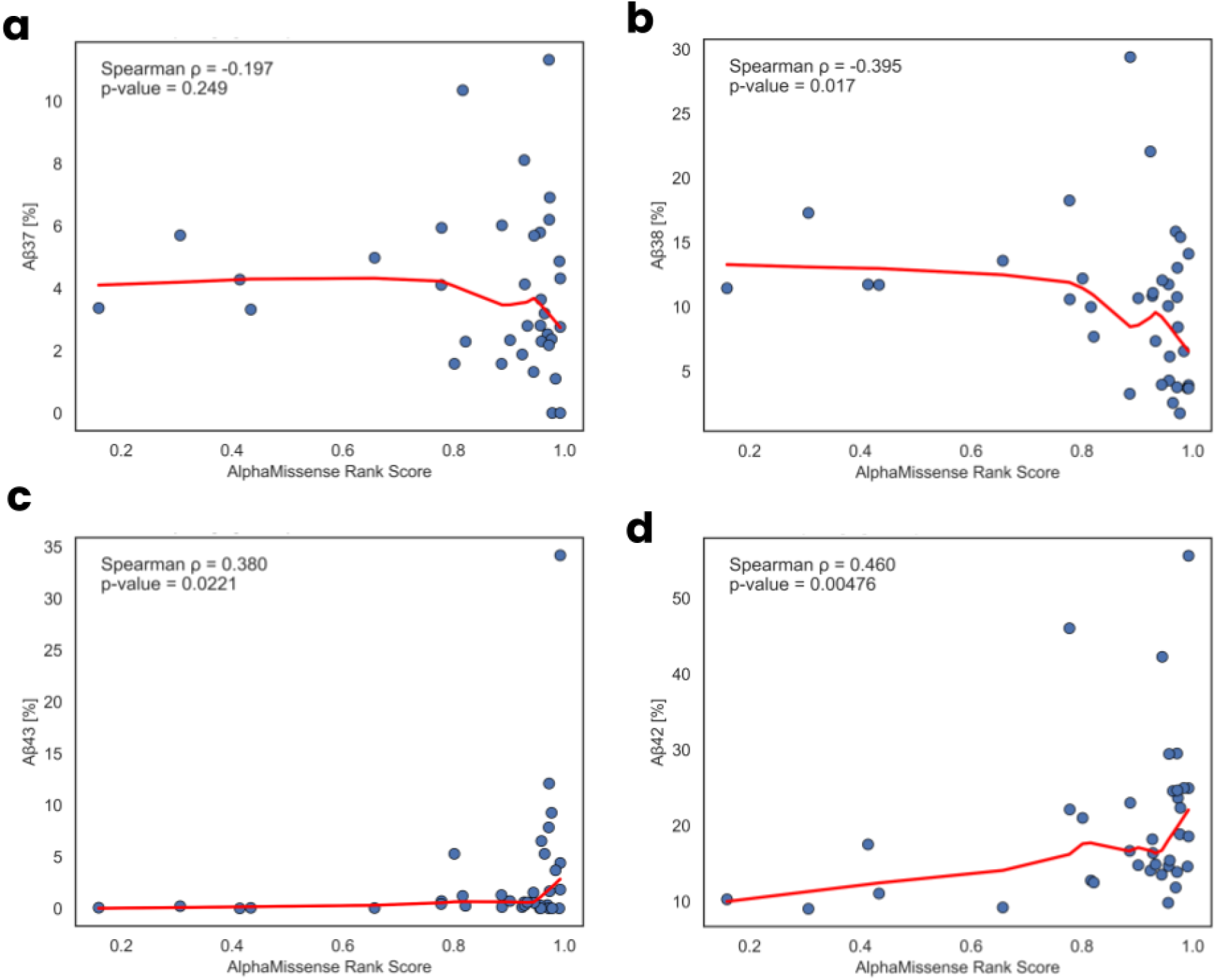
Correlations of AlphaMissense predictor with variants from Petit et al. (2022). **(a)** Correlation of Aβ37 levels with AM. **(b)** Correlation of Aβ38 levels with AM. **(c)** Correlation of Aβ43 levels with AM. **(d)** Correlation of Aβ42 levels with AM.

Lastly, we sought to resolve this controversy for the Aβ42/Aβ40 ratio using the sample from our previous study, Pillai et al., (2025), shown in **Figure S6**. As expected, the majority of predictors were significantly positively correlated with Aβ42/Aβ40 levels (34/37; 91.89%). From individual Aβ40 levels, we detected strongly negative correlation in most correlations (33/37; 89.19%) that is likely increasing the ratio. Likewise, Aβ42 levels were not significantly correlated for most VEPs (2/37; 5.41%).

Our analyses continued to *APP* variants from Pillai et al., (2025) **(Figure S5)**. We found that the Aβ42/Aβ40 ratio was significant (20/37; 54.05%). Unfortunately, we did not detect significant trends in the Aβ42 (*n* = 4/37; 10.81%) and Aβ40 levels (*n* = 1/37; 2.70%) that led to this strong correlation. This is likely owing to the limited sample size of *APP* variants (*n* = 25). Lastly, we evaluated the *PSEN2* from Pillai et al., (2025) **(Figure S4)**. The Aβ42/Aβ40 ratio was significantly positively correlated (18/37;48.65%), and was led by marked increases in Aβ42 (24/37; 64.86%) and insignificant Aβ40 levels (3/37; 8.11%). Surprisingly, the correlations of Aβ42/Aβ40 ratios in *PSEN2* align with the conclusions from Petit et al., (2022).

### 3.2. Biophysical analyses reveal missense variants associated with EOAD support both loss-of-function and dominant negative mechanism

In **Figure 3A**, we display biophysical structures of all three proteins superimposed to Cryo-EM structures, indicating near perfect similarity. We calculated the RSA of the wild-type residue for which mutations occur. We first identified that AAO is not correlated with surface exposure in *PSEN1* (ρ = 0.1278; *p* = 0.1735) **(Figure 3B)**. Conversely, as surface exposure increased, the total activity of the *PSEN1* increased significantly (ρ = 0.2148; *p* = 0.0157) **(Figure 3C)**. In **Table 1**, we report the summary of pairwise correlations between surface exposure of missense variants and Aβ levels. For Sun et al., (2016), we detected positive correlation between surface exposure and individual Aβ42, Aβ40 levels but marginally failed to detect the Aβ42/Aβ40 ratio. All variables from Petit et al., (2022) were insignificant with RSA. Similar to Sun et al., (2016), we identified significant correlations with individual Aβ40 levels, and likewise for negatively correlated Aβ42/Aβ40 ratio. Therefore, it appears that solvent accessibility of residues do not strongly correlate with functional effects among EOAD mutations.

**Table 1.** Song et al., (2026).

| Study | Variable | Gene | rho | p-value |
| --- | --- | --- | --- | --- |
| Sun et al., (2016) | AAO | *PSEN1* | 0.1278 | 0.1735 |
| **Sun et al., (2016)** | **Total Activity** | ***PSEN1*** | **0.2148** | **0.0157** |
| Sun et al., (2016) | Aβ42/Aβ40 | *PSEN1* | -0.1886 | 0.0686 |
| **Sun et al., (2016)** | **Aβ42** | ***PSEN1*** | **0.1934** | **0.0300** |
| **Sun et al., (2016)** | **Aβ40** | ***PSEN1*** | **0.1989** | **0.0255** |
| Petit et al., (2022) | Aβ37 | *PSEN1* | -0.1770 | 0.3016 |
| Petit et al., (2022) | Aβ38 | *PSEN1* | 0.2568 | 0.1306 |
| Petit et al., (2022) | Aβ40 | *PSEN1* | -0.0101 | 0.9533 |
| Petit et al., (2022) | Aβ42 | *PSEN1* | -0.1230 | 0.4748 |
| Petit et al., (2022) | Aβ43 | *PSEN1* | -0.2715 | 0.1092 |
| **Pillai et al., (2025)** | **Aβ42/Aβ40** | ***PSEN1*** | **-0.5692** | **0.0001** |
| Pillai et al., (2025) | Aβ42 | *PSEN1* | 0.0820 | 0.5479 |
| **Pillai et al., (2025)** | **Aβ40** | ***PSEN1*** | **0.4280** | **0.0010** |
| Pillai et al., (2025) | Aβ42/Aβ40 | *PSEN2* | -0.0574 | 0.6741 |
| Pillai et al., (2025) | Aβ42 | *PSEN2* | -0.0287 | 0.8336 |
| Pillai et al., (2025) | Aβ40 | *PSEN2* | 0.1000 | 0.4634 |
| Pillai et al., (2025) | Aβ42/Aβ40 | *APP* | 0.0300 | 0.8264 |
| Pillai et al., (2025) | Aβ42 | *APP* | 0.0011 | 0.9935 |
| Pillai et al., (2025) | Aβ40 | *APP* | -0.0024 | 0.9862 |

**Figure 3.**
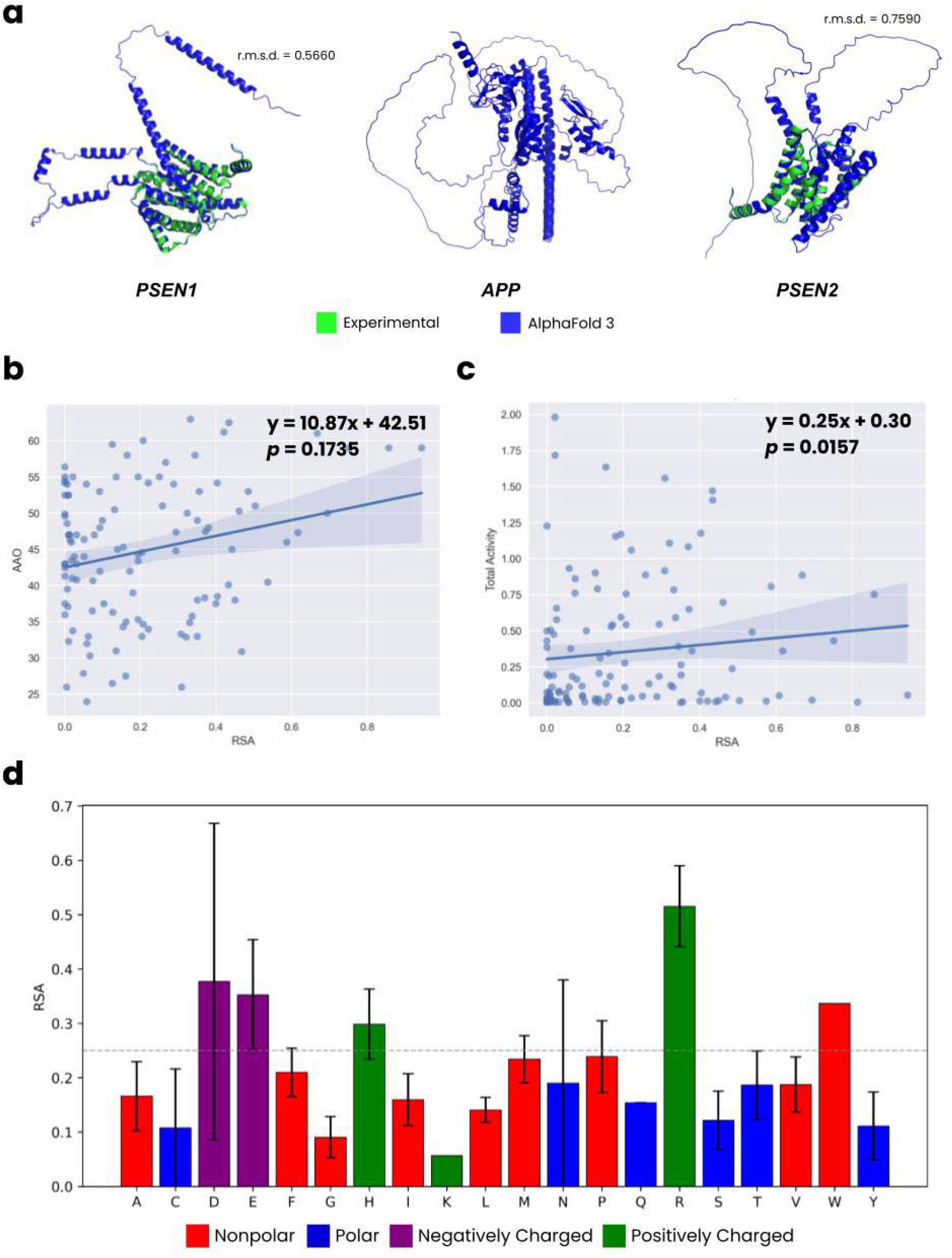
Biophysical characterization of pathogenic *PSEN1, APP*, and *PSEN2* variants. **(a)** Superimposition of AlphaFold 3 predictions of *PSEN1, PSEN2*, and *APP* over experimental CryoEM structures of PSEN1 and PSEN2 along with the room mean square deviation (r.m.s.d). **(b)** Non-significant positive correlation between RSA and AAO. **(c)** Correlation of RSA with total activity. **(d)** Distribution of RSA among wild-type residues of pathogenic PSEN1 variants. The dotted line indicates whether the protein is buried or exposed.

Finally, we sought to stratify the EOAD associated missense variants based on side-chain properties of the wild-type residue to further contextualize mutational effects. We aggregated all variants by gene, accounting for duplicate variants from independent samples, and showed the total count by RSA in **Figure 3D** for *PSEN1* and **Figure S6-7** for *APP* and *PSEN2*. We have set a threshold at 0.25 to indicate whether the residue type is buried or surface exposed. For *PSEN1*, we observe that mutations occurring on positively (Arg, His) and negatively (Asp, Glu) charged residues were generally exposed on the surface of the protein. By the same token, mutations occurring in hydrophobic residues were more buried in *PSEN1* with exception to tryptophan residue type.

Unfortunately, due to limited sample size for *PSEN2* and *APP*, this trend was correct for surface exposed charged amino acids but not for hydrophobic residues. It is broadly accepted that buried mutations cause a loss of stability, indicating a loss-of-function (LoF) mechanism (Cagiada et al., 2025). Likewise, dominant negative (DN) mechanisms act when the mutant interferes with the activity of a wild-type complex (Gerasimavicius et al., 2022). Therefore, our biophysical data shows mechanistic heterogeneity for EOAD. Charged surface variants preserve folding but likely alter protein-protein interactions, consistent with DN, and buried hydrophobic variants cause protein misfolding, consistent with LoF mechanism. It may be possible that there are multiple pathogenic mechanisms for EOAD.

## DISCUSSION

In this study, we performed extensive pairwise correlations of *in vitro* assays and patient data for pathogenic variants of *PSEN1, PSEN2*, and *APP* with VEPs and biophysical models. This study was motivated by the lack of diagnostic approaches to characterize variants in EOAD, leading to extensive analyses of VEPs. We make an important contribution to translating the landmark findings of Sun et al., (2016) and Petit et al., (2022) to identify broad clinical correlations for EOAD. We show consistently that VEP is correlated with increased Aβ42/Aβ40 levels. However, the cause for this correlation is debated among individual Aβ42 and Aβ40 levels. Our data also indicates that VEPs are well-correlated with AAO, a crucial predictor variable in precision medicine approaches for EOAD. Lastly, using full-length biophysical structures of each amyloidogenic gene, we show that missense variants display hallmarks of both DN and LoF effects to the complex. Therefore, this study offers wide-spread validation of VEPs and biophysical techniques for precision medicine applications for EOAD.

The potential cause(s) for increased Aβ42/Aβ40 levels is of particular controversyOur previous study (Pillai et al., 2025) and Sun et al., (2016) both agree that decreased Aβ40 levels caused the increased levels, but Petit et al., (2022) identified increased Aβ42 levels as the driving factor. Surprisingly, the data for *PSEN2* from Pillai et al., (2025) was in agreement with the conclusion from Petit et al., (2022). Petit et al., (2022) argued that the inconclusive findings from Sun et al., (2016) is owing to the experimental conditions with detergent extraction. Our pairwise correlations appear in agreement with Petit et al., (2022). However, further experimental studies are necessary to gain an ultimate understanding over where decreased Aβ40 or increased Aβ42 levels cause this important biomarker of EOAD.

Our study also shows the potential of VEPs to aid in clinical diagnostics among EOAD variants. Nevertheless, it remains to be determined whether VEPs have the ability to decipher mutation pathogenicity among variants with varying levels of pathogenic effects. In our data, though there is a negative logarithmic relation between predictors and AAO, it is important to note that a few variants are predicted as benign and occur at both earlier and late onset (∼40s, ∼60s) despite being pathogenic from Sun et al., (2016). Therefore, further studies are necessary to correlate VEPs and AAO. We also suggest pairwise correlations with longer Aβ strands, as suggested by Petit et al., (2022), which may be beneficial for molecular diagnostic approaches.

Lastly, we attempted to address the crucial question of the presenilin hypothesis: do pathogenic variants exhibit dominant negative effects (DN) structurally or a loss-of-function (LoF) mechanism? Structurally, there are pathogenic variants occurring on buried hydrophobic residues, indicative of LoF effects. However, there are a minority of variants located on the surface, specifically those occurring on arginine. It is possible that these surface variants alter signaling, demonstrating DN effects. Taken together, we argue that future studies should emphasize deciphering mechanistic heterogeneity for EOAD.

## Data Availability

The data is available at https://github.com/jiyeonsongg/am_analysis.

## Acknowledgement

None to declare.

## Artificial Intelligence

None to declare.

## Consent Statement

The patient data used in this study is publicly available from Sun et al., (2016).

