## Supplementary Material for "Correlations of 276 missense variants of *PSEN1, PSEN2*, and *APP* on the production of Aβ peptides against variant effect predictors and biophysical structures"

Pre-print server: bioRxiv, CC-BY-NC-ND 4.0.

Classification: Research Article, XXXX words

Keywords: Alzheimer’s disease; AlphaMissense; Missense Variant Effect Predictors.

**Figure Description**

**Figure S1.** Heatmap of variant effect predictors correlated against the clinical and biochemical variables from Petit et al., (2022).

**Figure S2.** Heatmap of variant effect predictors correlated against the clinical and biochemical variables from Pillai et al., (2025).

**Figure S3.** Heatmap of variant effect predictors correlated against the clinical and biochemical variables from Sun et al., (2016).

**Figure S4.** Heatmap of variant effect predictors correlated against the clinical and biochemical variables from Pillai et al., (2025).

**Figure S5.** Heatmap of variant effect predictors correlated against the clinical and biochemical variables from Pillai et al., (2025).

**Figure S6.** Total EOAD missense variant gene count according to side-chains using RSA for APP from Song et. al., (2026).

**Figure S7.** Total EOAD missense variant gene count according to side-chains using RSA for PSEN2 from Song et. al., (2026).

Figure S1. Song et al., (2026).

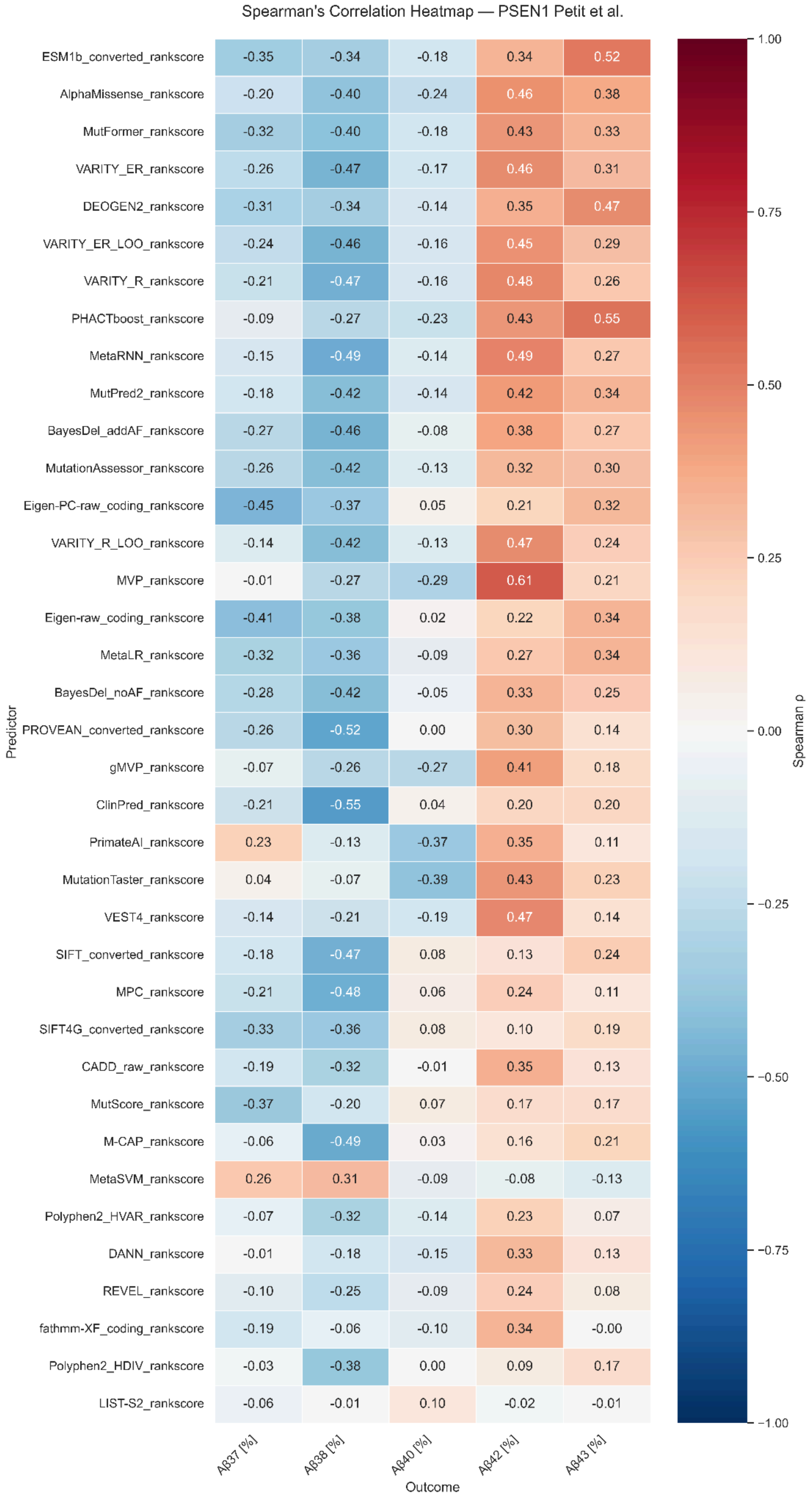

Figure S2. Song et al., (2026).

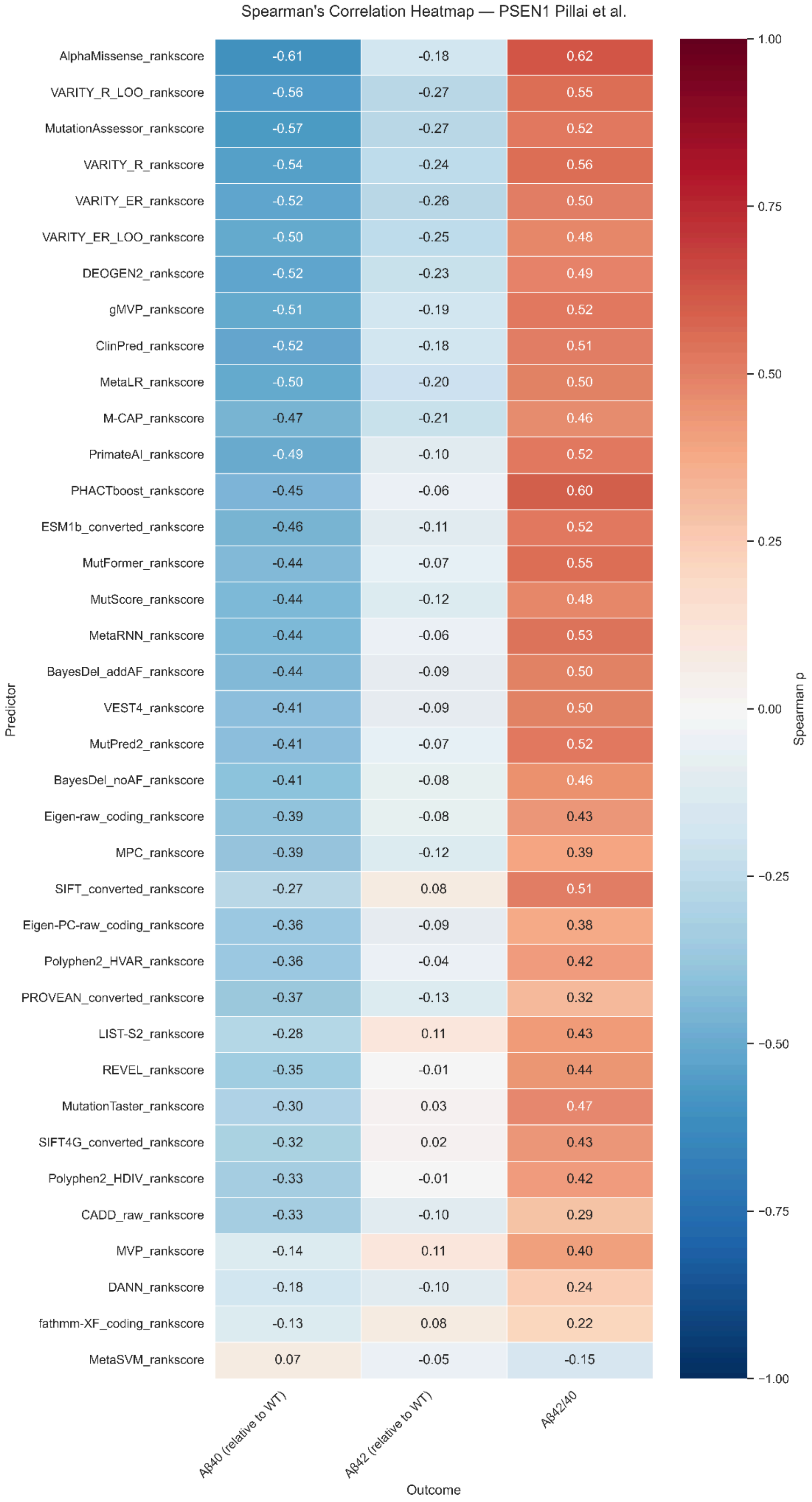

Figure S3. Song et al., (2026).

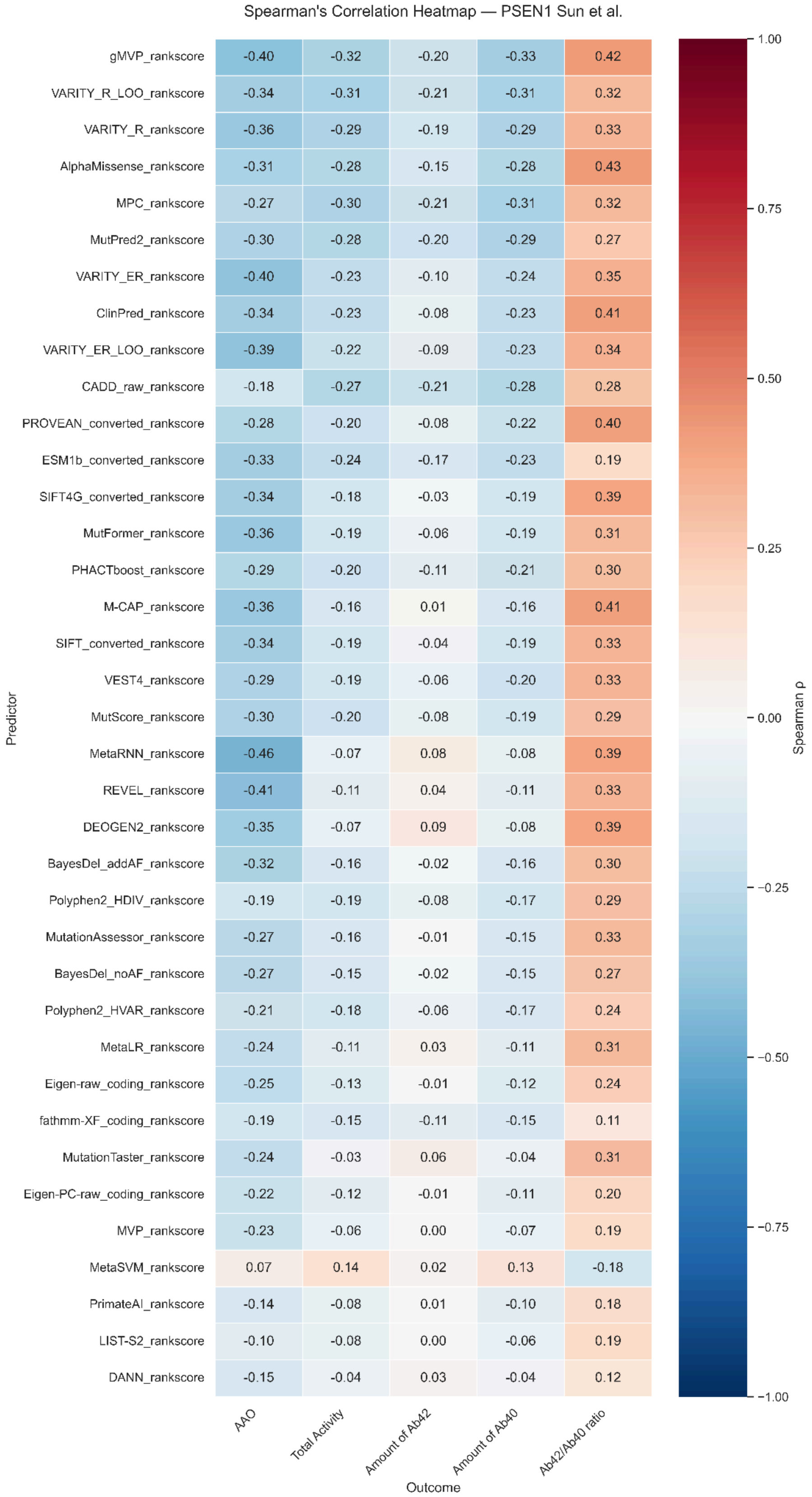

Figure S4. Song et al., (2026).

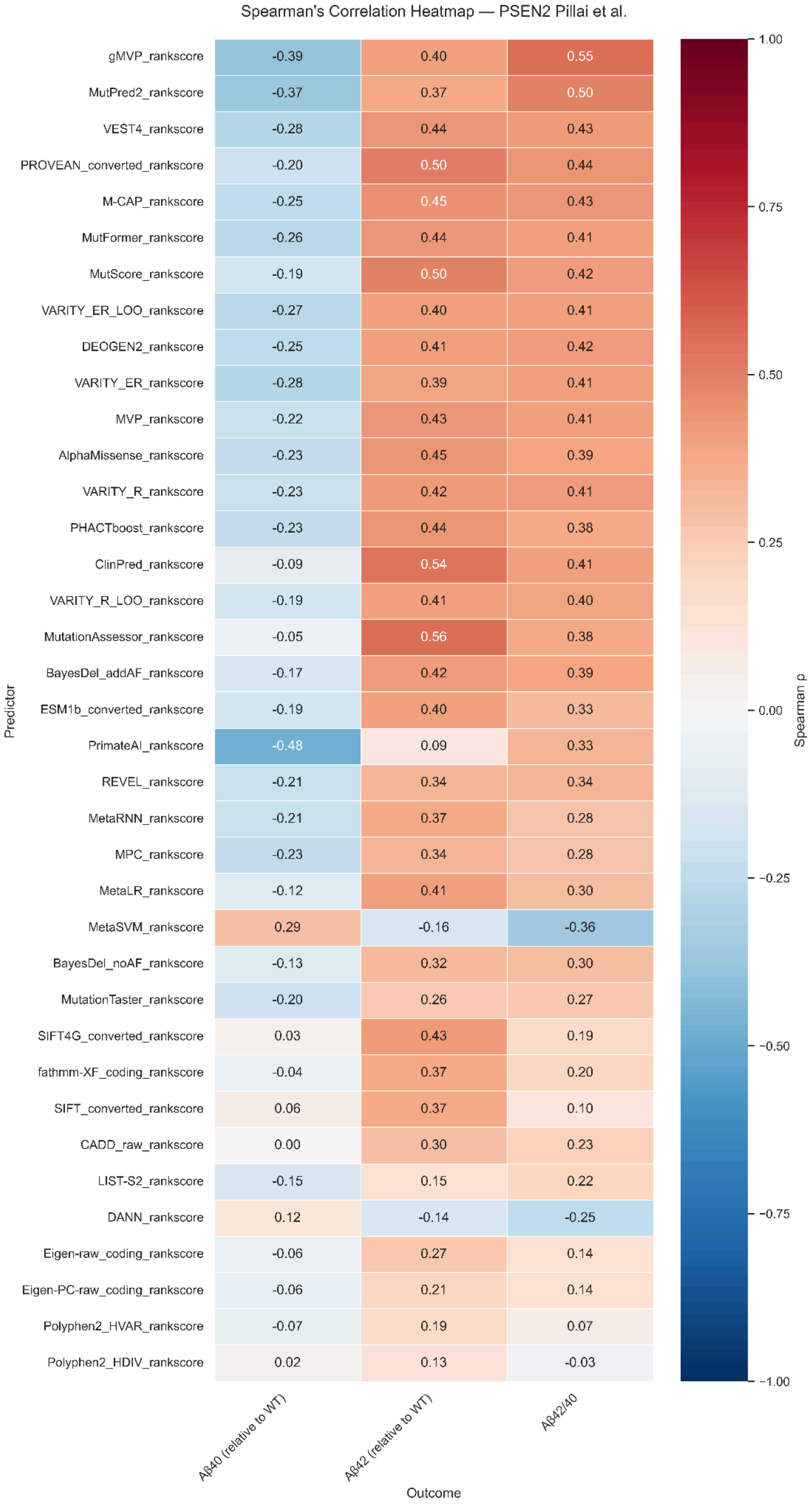

Figure S5. Song et al., (2026).

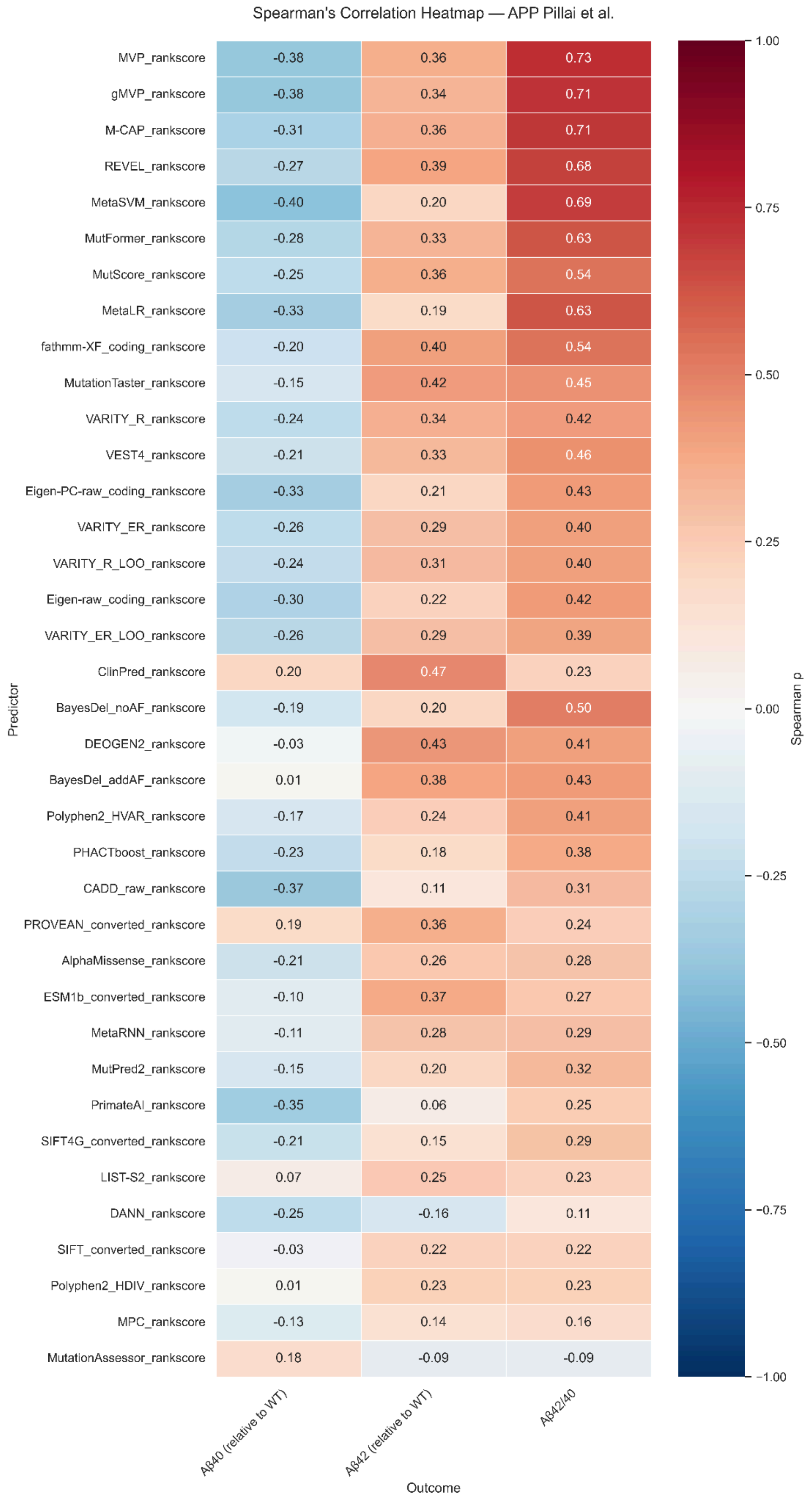

Figure S6. Song et al., (2026).

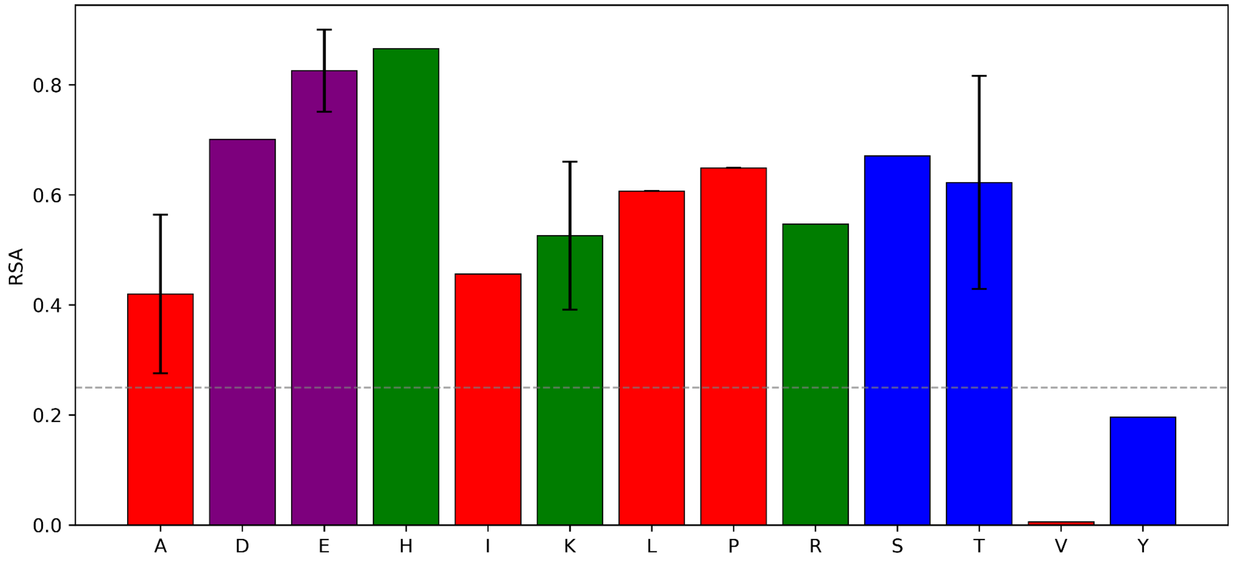

Figure S7. Song et al., (2026).

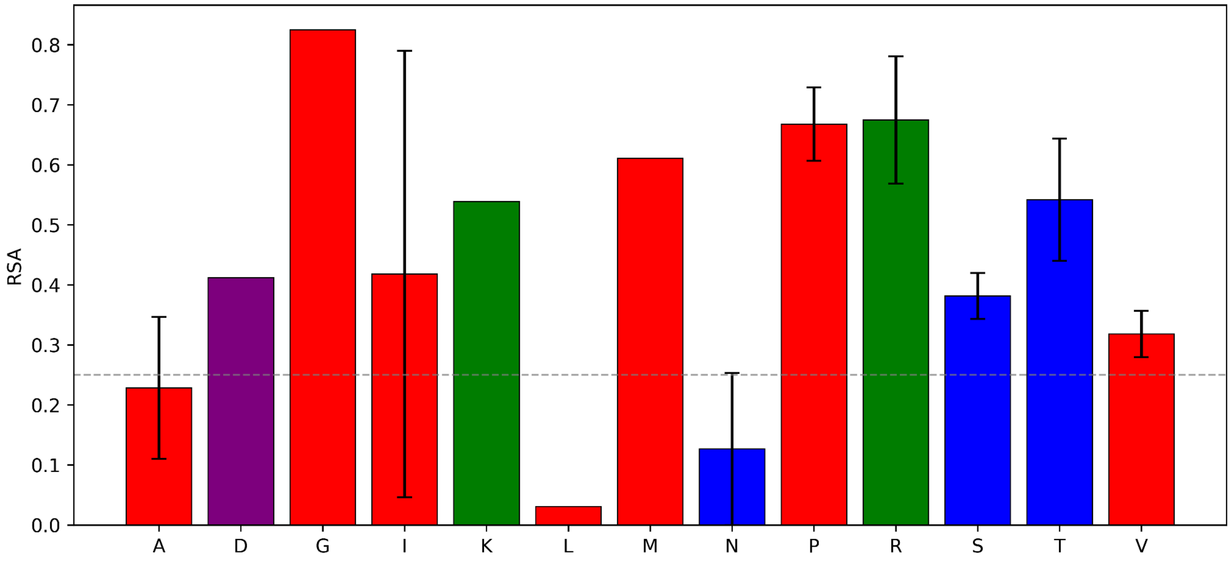
